# The catabolic nature of fermentative substrates influence proteomic rewiring in *Escherichia coli* under anoxic growth

**DOI:** 10.1101/2024.09.19.613855

**Authors:** Huda Momin, Deepti Appukuttan, K.V Venkatesh

## Abstract

During anaerobic batch fermentation by *Escherichia coli*, there is a decline in cell proliferation rates and a huge demand is placed on cellular proteome to cater its catabolic and anabolic needs under anoxic growth. Previous studies have established a direct relationship between *E. coli* growth rate and cellular ribosomal content for fast proliferating cells. In this study, we integrated experimental findings with a systemic coarse-grained model of proteome allocation, to characterize the physiological outcomes at slow growth rate during anaerobic fermentative catabolism of different glycolytic and non-glycolytic substrates. The anaerobic catabolism of substrates favored high ribosomal abundances at lower growth rates. Interestingly, a modification of previously discussed “growth law”, the ratio of active to inactive ribosomal proteome was found to be linearly related to growth rate for cells proliferating in slow to moderate regime (growth rate < 0.8 h^-1^). Also, under nutrient- and oxygen-limited growth conditions, the proteome proportion allocated for ribosomal activity was reduced, and resources were channelized towards catabolic and metabolic activities to overcome the limitations imposed while uptake and metabolizing substrate. The energy intensive uptake mechanism or lower substrate affinity, expended more catabolic proteome, which reduced its availability to other cellular functions. Conclusively, the nature of catabolic substrates imposed either uptake limitation or metabolic limitation coupled with ribosomal limitation (arising due to anoxic and nutritional stress), which resulted in higher proteome expenditure leading to sub-optimal phenotype.

## Introduction

In a single-cell organism like *Escherichia coli*, the strategic allocation of cellular resources rewires physiological outcomes, fundamentally measured as cell proliferation rate under different growth conditions^1–4^. Also, the nutritional content of the growth medium influences cellular gene expression level and its macromolecular composition by orchestrating its growth rate^4–6^. The native host metabolism uniquely metabolizes incoming substrates, for the production of anabolic precursors and energy; thereby leading to polymerization of cellular macromolecules like protein, RNA, cell envelop, etc., in cellular machinery^7^. Within a fixed cellular budget, a bacterial cell balances the production of proteins and their synthesizing machinery called ribosomal assembly^3^. Altogether, the cellular ribosomes are critical in shaping its protein repertoire and the level of gene expression under different growth conditions^4,8^. The cellular ribosome abundance is reflected by the total RNA-to-protein (R/P) ratio, as *E. coli* rRNA (∼ 85% of total RNA) is folded in ribosomes^4,9^ and > 95% of total RNA (including rRNAs and tRNAs) deals with protein translation^3,5^. Moreover, *E. coli* ribosomes exhibit distinctive mass composition, of RNA and protein in a ratio 2:1, unique for cell growth maximization^10,11^.

The empirical linear relationship between ribosomal abundance and growth rate for *E. coli* cells proliferating in fast to moderately slow regime (doubling time ∼ 20 min to 2 hours)^4,5,12,13^ has been physiologically defined as the “growth laws”^4,9^. In a batch culture with limited ribosomes translating at a constant rate, the *E. coli* growth rate can be adjusted by modulating medium composition, such as by adding supplements like amino acids, vitamins, and nucleosides, or altering the carbon source, which favor shorter doubling time and faster proliferation rates. Conversely, when the ribosomal translation rate is impaired by antibiotics in a graded manner in a medium with fixed nutrient composition, there is a decline in growth rate with an increase in ribosomal abundance^4,8^. This relationship can be improved by enhancing the nutrient medium quality. In both the cases the nutrient medium quality can be critical in defining the physiological outcome. Furthermore, integrating these key observations on ribosome-growth rate relationships with coarse-grained proteome partitioning^4^ based on functionality, enhances our understanding of *E. coli* physiology. The studies including specific protein abundance map and system-wide proteome allocation under different growth conditions have been previously reported^14–16^. The ubiquitous microbial phenomenon such as overflow metabolism^17^ or carbon catabolite repression^1^ has been well explained by considering the cost of protein synthesis. The knowledge of cellular proteomes has been integrated with a systemic approach to uncover the physiological variations of *E. coli*^18–21^. However, there exist a knowledge gap regarding the resource rewiring in *E. coli* during anaerobic fermentative substrate catabolism.

The anaerobic fermentation is an independent nutritional mode exhibited by *E. coli* cells for the catabolism of different substrates. Although the mode of nutrition has certain limitations in terms of energy biogenesis, substrate oxidation, carbon wastage, etc., it is an appealing mode for the production of various chemicals of industrial significance. So, here we are interested in understanding the role of ribosome in determining bacterial growth rate during anaerobic fermentative breakdown of different substrates. We also studied the rewiring of proteome allocation with changing substrates and growth rate during anaerobic batch fermentation. Here, we considered six substrates viz., glucose, fructose, xylose, sorbitol, gluconate and pyruvate, which varied in their chemical nature including carbon content (3, 5 or 6 carbon), catabolic pathways (glycolytic or non-glycolytic), and oxidation state (−1, 0, +1). For understanding the influence of substrate in proteome allocation in *E. coli*, we cultivated cells in minimal growth medium M9 supplemented with MOPS buffer, to rule out the interventions of growth medium in physiological outcome. We resorted to theoretical simulations by constraining the experimental results to generate a coarse-grained model of proteome allocation. Thus, we elucidated the role of ribosomes and proteome partitioning in defining the *E. coli* physiology during anaerobic fermentation of different substrates.

## Materials and methods

### The bacterial strain and its growth condition

The *Escherichia coli* K-12 strain BW25113 [△*(araD-araB)567* △*(rhaD-rhaB)568* △*lacZ4787* (::rrnB-3) *hsdR514 rph-1*]^22^ was used for anaerobic fermentation of different substrates. The bacterial cells were anaerobically cultured at 37°C, with rotor speed at 120 rpm, in 200 mL sterile M9 minimal medium (composition per litre: 6 g anhydrous Na_2_HPO_4_, 3 g KH_2_PO_4_, 0.5 g NaCl, 1 g NH_4_Cl, 1 M MgSO_4_, 1 M CaCl_2_) supplemented with 100 mM MOPS (3-(N-morpholino) propane sulfonic acid) as a buffering agent. The pH of the medium was set to 7.2 and the anaerobic condition was maintained by sparging sterile nitrogen gas in the reaction assembly. The six substrates viz., glucose (2 g L^−1^), fructose (2 g L^−1^), xylose (2 gL^−1^), sorbitol (2 g L^−1^), gluconate (2.4 g L^−1^), and pyruvate (2.4 g L^−1^) were added such that equal amount of carbon was maintained under different growth conditions. The chemicals used in this study were purchased from Merck. The bacterial growth rate was determined spectrophotometrically (Thermo Scientific Multiscan GO) by measuring cellular density at 600 nm (O.D_600_).

### Total RNA estimation by TRIzol method

Total RNA quantification from three biological replicates for each substrate was done by the TRIzol method^23^ with slight modifications. The bacterial cells were harvested from mid-exponential phase culture of anaerobic fermentation with an optical density of 20 by centrifuging it at 7000 rpm, 4°C for 20 mins. The cells pellet was washed with M9 + MOPS medium, snap-frozen in liquid nitrogen and stored in a -80°C freezer until RNA extraction. For extracting total RNA, the cells were suspended in 250 µL TRIzol reagent and were homogenized with a sterile micropestle for 1.5 mins, this process was repeated twice. Additionally, 500 µL TRIzol reagent was introduced in the centrifuge tube and the suspension was homogenized by vortexing briefly (∼15-30 secs), followed by incubation at room temperature for 5-7 mins for complete dissociation of nucleoproteins complex. Next, 300 µL of chilled chloroform was added to the tube, mixed thoroughly by shaking, and incubated for 5 mins. The samples were then centrifuged at 13500 rpm, 4°C for 20 mins. The upper aqueous phase was carefully transferred into a fresh micro-centrifuge tube and 800 µL of chilled isopropanol was added to it. It was then incubated at -20°C overnight. The next day, the samples were centrifuged at 13500 rpm, 4°C for 30 mins, forming a white pellet. This pellet was washed twice with 1 mL 75 % chilled ethanol. The pellet was air-dried until translucent. The resulting pellet was suspended in 20 µl nuclease-free water and incubated at a 55°C dry bath for 10-13 mins. The concentration along with purity was checked using nanodrop and the quality check was done by agarose gel electrophoresis. Any genomic DNA contamination in the extracted total RNA was removed by DNase treatment, followed by phenol-chloroform extraction.

### Total protein estimation by Biuret test

For the Biuret method^1^ of total protein quantification, cells from mid-exponential phase culture with an optical density of 2 were collected by centrifuging at 7500 rpm, 4°C, 20 mins.The resulting pellet was washed with autoclaved Milli Q and resuspended in 200 µL water. It was fast – frozen in liquid nitrogen and thawed in water bath at room temperature for 5 mins. 100 µL of 3 M NaOH was added to the samples and it was incubated at 100°C heat block for 5 mins, for protein hydrolysis. The resulting samples were cooled in a water bath at room temperature for 5 mins. Later, 100 µL of 1.6 % CuSO_4_ solution was added, followed by thorough mixing at room temperature for 5 mins. Finally, the samples were centrifuged and the absorbance was measured at 555 nm. The protein concentration in the sample was quantified from the BSA standard curve, prepared by the same biuret reaction.

### Genome-scale Metabolic Model (GEM) and Constrained Allocation Flux Balance Analysis (CAFBA)

The genome-scale metabolic model *i*JO1366^24^ for *E. coli* MG1655 (*E. coli* K-12 family) was used for all simulations with certain modifications unique to the *E. coli* BW25113 strain. The *E. coli* strain BW25113 differs from MG1655 with several gene deletions like *lacZ, araBAD*, and *rhaBAD*. Consequently, the fluxes across reactions like LACZ, RMK, RMI, RBK_L1, RMPA, LYXI, and ARAI were constrained to zero. Moreover, the oxygen exchange reaction was constrained to zero, for mimicking the anaerobic fermentative growth. Constrained Allocation Flux Balance Analysis^19^ was used to systematically allocate *E. coli* proteome resources into coarse-grained functional sectors. These sectors comprise genes of common interests such as substrate catabolism, anabolism, housekeeping, and maintenance^1,4^, etc. The CAFBA simulations were performed in MATLAB (The MathWorks Inc., MA, USA) using Gurobi Solver (Gurobi Optimization, USA). For our nutrient-limited study, we have systematically categorized the *E. coli* BW25113 proteome into four coarse – grained sectors as:

1. Ribosomal (R) sector (∅_*R*_): This class comprises ribosomal and its affiliated proteins.
2. Catabolic (C) sector (∅_*C*_): It includes substrate influx and transport proteins.
3. Metabolic (M) sector (∅_*M*_): This class accommodates metabolic enzymes.
4. Housekeeping (Q) sector (∅_*Q*_): It comprises the growth-independent core proteome.

### Statistical Analysis

The results reported here are mean of three independent biological replicates (n=3), represented along standard deviations. The Student’s T-test was performed to determine the statistical significance across different growth conditions. Since the experiments were carried out in triplicates, simulations were run thrice for each case to minimize theoretically associated errors. The corresponding deviations are reported along with actual values.

## Results

### The inactive ribosomal abundance influenced total protein content across different growth conditions

The cellular ribosomal abundance determines its protein content. We measured cellular RNA-to-protein ratio (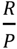ratio, r) or ribosomal abundance during anaerobic fermentation of glucose, fructose,xylose, sorbitol, gluconate, and pyruvate. However, there was no significant difference in ribosomal abundance during the fermentative breakdown of these substrates (Fig. 1 (A)). Contrarily, total cellular protein content significantly varied across substrates (except glucose and fructose had similar protein content) (Fig. 1 (B)). The substrates like xylose, sorbitol, gluconate, and pyruvate had total protein content comparatively lesser than glucose and fructose. Since ribosomes are protein-synthesizing machinery, we were intrigued by the non-significant difference in ribosomal abundance but a significant protein variation during the anaerobic fermentative catabolism of different substrates.

**Figure 1.**
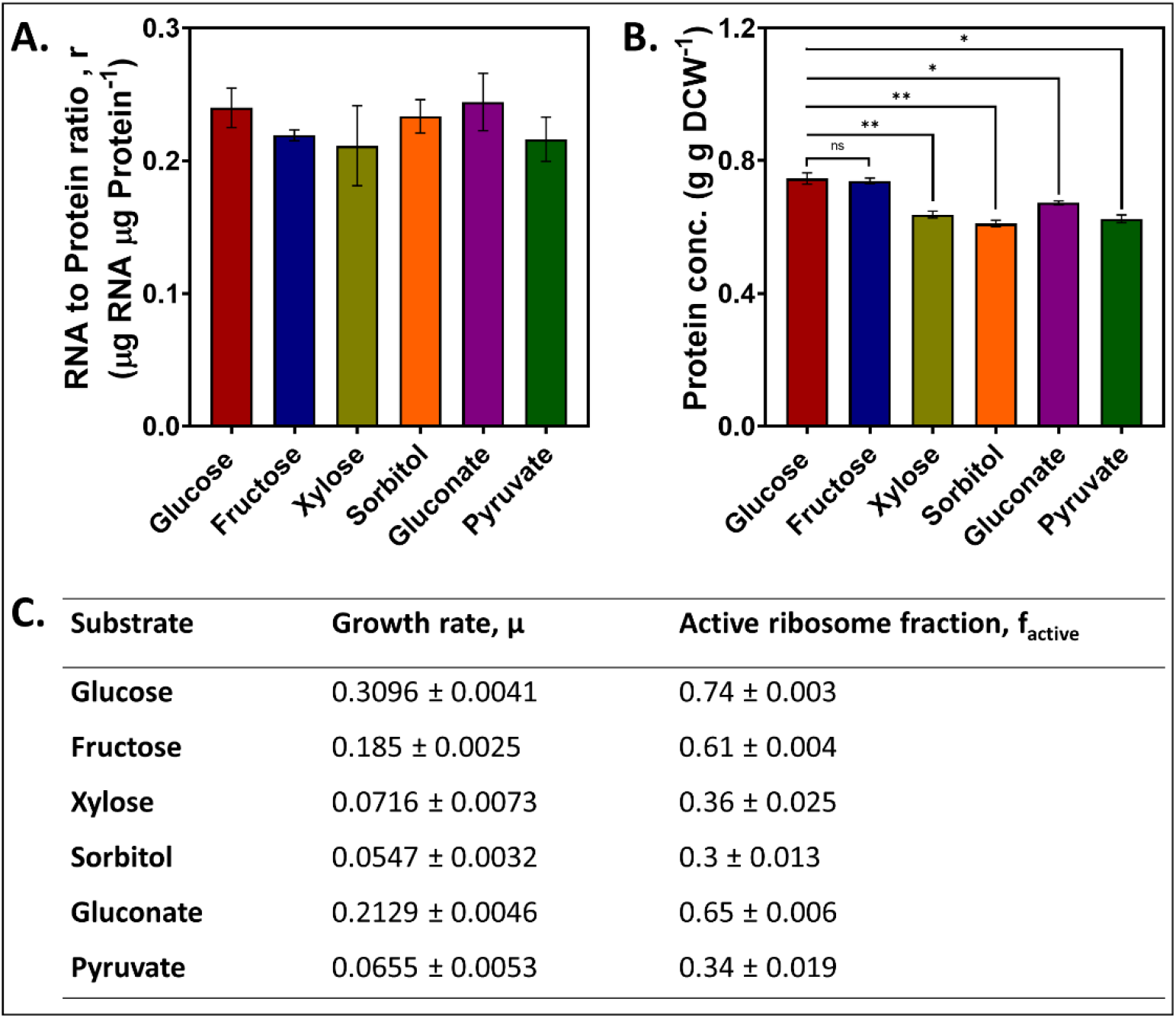
(A) The 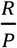 ratio during anaerobic batch fermentation of different glycolytic and non-glycolytic substrates. (B) The total protein content during anaerobic growth of *E. coli* BW25113 on different substrates. The asterisks represent the level of significance as p-value < 0.05 (Student’s t-test); ns: non-significant. (C) The theoretically calculated active ribosome fraction in *E. coli* BW25113 during anaerobic catabolism of different substrates.

Also, *E. coli* cells exhibited unique specific growth rates in these culture conditions. The bacterial ribosomes play a key role in shaping its proliferation rate. Previously, a decline in the active ribosomal fraction was correlated with growth rate retardation^25^ and we observed a Michaelis-Menten type of relationship between active ribosomal fraction and *E. coli* growth rate (Supplementary, S.1). As a result, we calculated the active ribosomal fraction which was responsible for different *E. coli* proliferation rates during anaerobic catabolism of glycose, fructose, xylose, sorbitol, gluconate and pyruvate respectively (Fig. 1 (C)). At lower cell proliferation rates, the fraction of actively translating ribosomes declined, reducing cellular protein content. Also, we theoretically determined the ribosomal translation elongation rate (*k*, amino acid per second) by using the formula 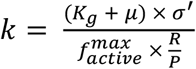 it increased with growth rate for maintaining the protein pool required for cellular maintenance and proliferation under nutrient-limited anaerobic growth (Fig. S1 (B)). Thus, the physiological outcome was a consequence of an interplay between active cellular ribosome content and translational elongation rate that eventually influenced bacterial cell proliferation rate.

### The ratio of active to inactive ribosomal proteome determined the cell proliferation rate

The cellular ribosomes as a protein translating entity, themselves require proteins (ribosomal and its affiliated proteins) for functionality. Consequently, the protein repertoire available for other cellular activity reduces. As ribosome-affiliated proteome fraction is directly linked to cellular ribosomal abundance (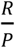ratio) (Supplementary, S.2), the total ribosomal proteome share was similar during the metabolism of all six substrates (Fig. 2 (A)). However, active ribosomal proteome had the highest share during glucose catabolism, followed by gluconate and fructose metabolism. During the breakdown of xylose, sorbitol, and pyruvate, the inactive ribosomal proteome share increased, thereby imparting a retarded growth physiology.

**Figure 2.**
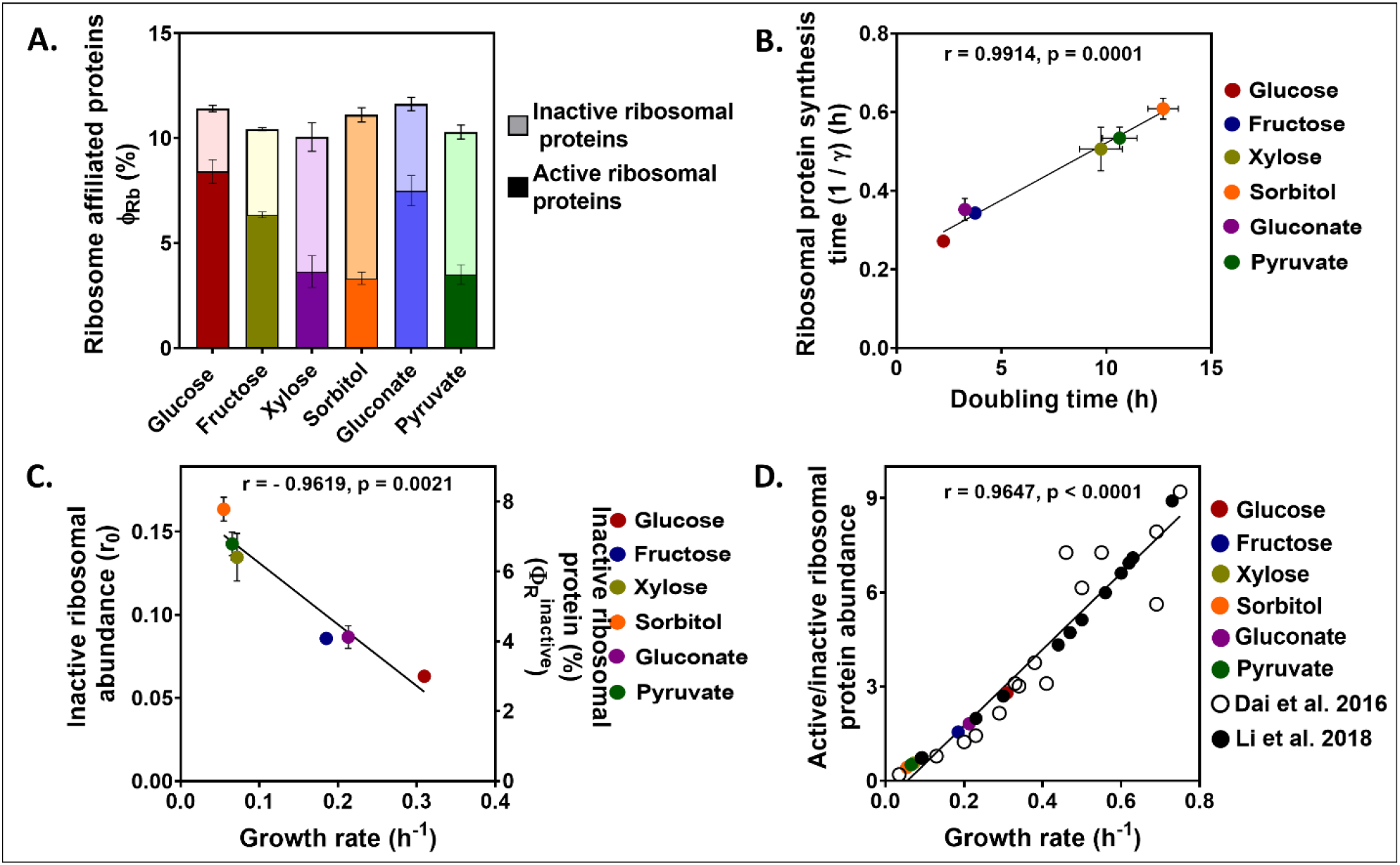
The role of the ribosomal-affiliated proteome in determining cell growth rate during the anaerobic fermentative breakdown of different glycolytic and non-glycolytic substrates. (A) The relative proportion of active and inactive ribosomal proteome share. (B) The influence of ribosomal protein synthesis time on cell doubling time. (C) The influence of inactive ribosomes and their affiliated proteome on cellular growth rate. (D) The relationship between ratio of active-to-inactive ribosomal proteome abundance and cellular growth rate as obtained from this study and those calculated from literature data^3,25^. The Pearson’s ‘r’ and ‘p’ values indicate the level of significance; r > 0.8 and p < 0.075.

The ribosomal protein synthesis time varied with cell doubling time (Fig. 2 (B)). Thus, longer the time required to synthesize ribosomal proteins, the slower was the bacterial growth rate. During catabolism of substrates like xylose, pyruvate and sorbitol, the actively translating ribosomes became limiting, which increased the ribosomal protein synthesis time, eventually leading to growth retardation. As a result, the ribosomal protein synthesis time was a crucial determinant of *E. coli* BW25113 physiology during the anaerobic fermentative breakdown of glucose, fructose, xylose, sorbitol, gluconate, and pyruvate. Moreover, we observed a negative influence of inactive ribosomal abundance, along with inactive ribosomal proteome share on bacterial growth rate during anaerobic fermentative breakdown of different glycolytic and non-glycolytic substrates (Fig. 2 (C)). Thus, under nutrient limited growth, a slow *E. coli* BW25113 growth rate can be attributed to a larger proportion of inactive ribosomal proteome. This can be the outcome of amino acid scarcity resulting from substrate catabolism in a minimal growth medium. Additionally, an alternative to linear growth laws established previously^4,25^, we observed a direct influence of fractional active to inactive ribosomal proteome ratio on bacterial growth rate. This observation was supported by data consolidated from literature for *E. coli* cells^3,25^ proliferating in slow to medium growth regime (growth rate < 1 h^-1^) (Fig. 2 (D)). Alternatively, this correlation even holds true for the fractional ratio of active to inactive ribosomes for cells proliferating in the same regime. This is an interesting outcome of the study as it determines the extent to which ribosomes and its affiliated proteome can influence *E. coli* growth rate under nutrient-limited growth conditions.

### The cost associated with catabolic sector proteome was determined by the substrate uptake mechanism and catabolism

For determining the proteome share associated with substrate influx during their anaerobic fermentative breakdown, we first calculated the associated proteome cost for the influx of each substrate. For each simulation, the experimentally determined metabolite secretion rates under anaerobic growth condition was considered along with the parameters enlisted in supplementary, S.3. The experimentally determined substrate uptake rate was kept unbound while optimizing the growth rate. The catabolic proteome cost was determined by iterating value ranges 0 and 1 (100 iterations). The (*w*_*C*_) value corresponding to predicted substrate uptake rate and growth rate, that matched with experimentally determined values were considered. The study involved equal number of carbon during anaerobic fermentation of all six substrates. As a result, corresponding proportion of substrates varied, in order to compensate the required carbon content in growth medium. Depending upon variable substrate affinity and its uptake mechanism, the substrate influx rate and bacterial growth rate varied.

The bacterial cells had least hindrance in pyruvate uptake (least uptake cost), owing to dual uptake mechanism^26–28^ operative in *E. coli* cell. However, its non-glycolytic catabolism failed to cope with optimal performance, giving slower proliferation rates. Next in line, was glucose internalization, followed by gluconate influx-associated cost. The substrate internalization rates for both the substrate were similar, however, variation existed in bacterial growth rates due to different metabolic pathways. The xylose-associated influx cost was comparatively lower than fructose and sorbitol internalization. Although the uptake rate was sufficient, the bacterial proliferation rate was compromised here due to energy intensive substrate internalization^29^. The fructose associated cost was at higher side, however it supported decent anaerobic bacterial growth rate. Lastly, sorbitol influx was a draining task for *E. coli* BW25113 cells that led to lowest bacterial growth rate. Thus, theoretically calculated catabolism-associated proteome cost was an outcome of limiting substrate behavior comprising of substrate uptake rate, the substrate metabolic pathway, the cell proliferation rate, etc. (Fig. 3 (A)).

**Figure 3.**
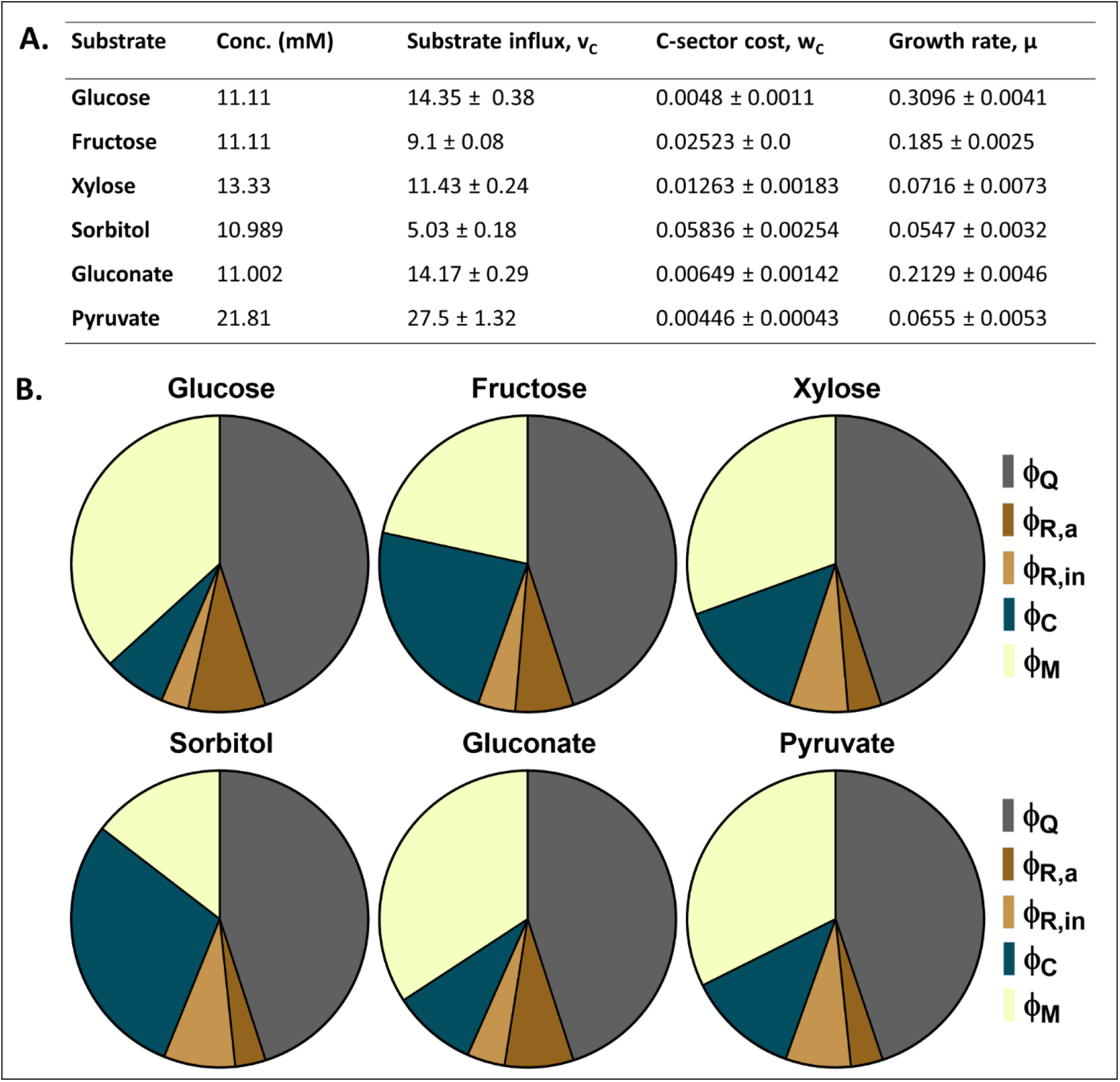
(A) Theoretically calculated C-sector proteome cost in *E. coli* BW25113. (B) The proteome partitioning to different cellular functions during anaerobic batch fermentation of different substrates. The proteome fractions are named as: ∅_*Q*_ -housekeeping (Q) sector, ∅_*R,a*_ -active ribosomal (R) sector, ∅_*R,in*_ - inactive ribosomal (R) sector, ∅_*C*_ - catabolic (C) sector, and ∅_*M*_ -metabolic (M) sector.

### The nature of substrate influenced coarse-grained partitioning of E. coli proteome during anaerobic fermentative breakdown

The coarse-grained allocation of proteome into four functional sectors was done to understand how cellular resource partitioning shape bacterial physiology. In *E. coli*, the core proteome share is independent of bacterial proliferation rate and constitute 45%^4^ of total cellular proteome. The ribosomal proteome share was jointly calculated by experimentation and theoretical considerations. The relative active and inactive ribosomal share is depicted alongside the housekeeping proteome sector (Fig. 3 (B)). Using the constrained allocation flux balance analysis, we were able to quantify the proteome cost associated with nutrient influx, thus the catabolic sector proteome was determined. Lastly, the metabolic sector proteome was grouped as remaining cellular proteome and calculated as:

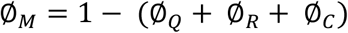

The driving flux for catabolic sector proteome was substrate influx rate, where relative distribution of C-sector proteome varied with type of substrate. The pyruvate influx happened at highest rate with least influx cost. However, the C-sector proteome occupied sufficient share (∼12%) of cellular proteome during pyruvate catabolism. The substrates, glucose and gluconate were internalized at similar rate, but their uptake mechanisms posed a difference in their catabolic proteome cost. This yielded different cell proliferation rates, and slightly higher proteome was allocated for gluconate (∼9%) influx as compared to glucose (∼7%) uptake. For xylose internalization, ∼15% proteome was channelized for its influx, whereas ∼23% proteome was utilized for fructose influx. The highest catabolic proteome share was compromised for sorbitol influx (∼29%). The catabolic proteome allocation for uptake of different substrates was resultant of tuning between substrate affinity to *E. coli* cell, and its uptake mechanism.

Once the catabolic proteome resumed their activity, the remaining biosynthetic proteome also become functional for carrying out metabolic conversions, leading to cell growth and multiplication. A substantial allocation of proteome towards substrate uptake would render lesser proteome for cellular biosynthesis. As a result, the sorbitol metabolism had least metabolic proteome (∼15%) share, proving insufficient for supporting decent bacterial growth rate. The glucose (∼37%) and gluconate (∼34%) had comparatively higher biosynthetic proteome leading to higher cell proliferation rates. Next in line, was pyruvate biosynthetic proteome (∼33%) capable of facilitating higher cell proliferation rate. However, the metabolic pathway breakdown of pyruvate rendered insufficient energy biogenesis and redox level within cell, that eventually compromised its growth rate. Moreover, similar was the case with xylose metabolism, where ∼ 31% of biosynthetic proteome was involved in shaping the bacterial physiology. However, the higher ATP requirement during substrate influx, compromised the ATP availability for other cellular functions, thereby lowering bacterial growth rate. Lastly, in spite of having comparatively lesser metabolic proteome, the fructose (∼22%) metabolism was adequate enough to support a decent bacterial growth.

### The absolute quantification of proteins belonging to different coarse-grained sectors were correlated with physiological parameters

The housekeeping proteome occupy a fixed fraction in cellular proteome resources, invariant to bacterial growth rate^4,9^. This sector constitute 0.45 fraction of *E. coli* proteome^4^. Upon quantifying the protein content of this sector, we found it to be linearly correlated with cell growth rate (Fig. 4 (A)). Thus, core proteome mass (by weight) responsible for cellular integrity and maintenance increased with cell proliferation rate. The cell biomass yield with respect to substrate uptake rate was also linearly related with core protein mass (Fig. 4 (B)). This implies the influence of substrate influx rate on core proteome mass. The higher ratio of cell growth to substrate, higher is the core proteome mass. Thus, core cellular proteome is influenced by cell proliferation rate, wherein substrate influx and its catabolism play a vital role in determining the phenotypic outcome.

**Figure 4.**
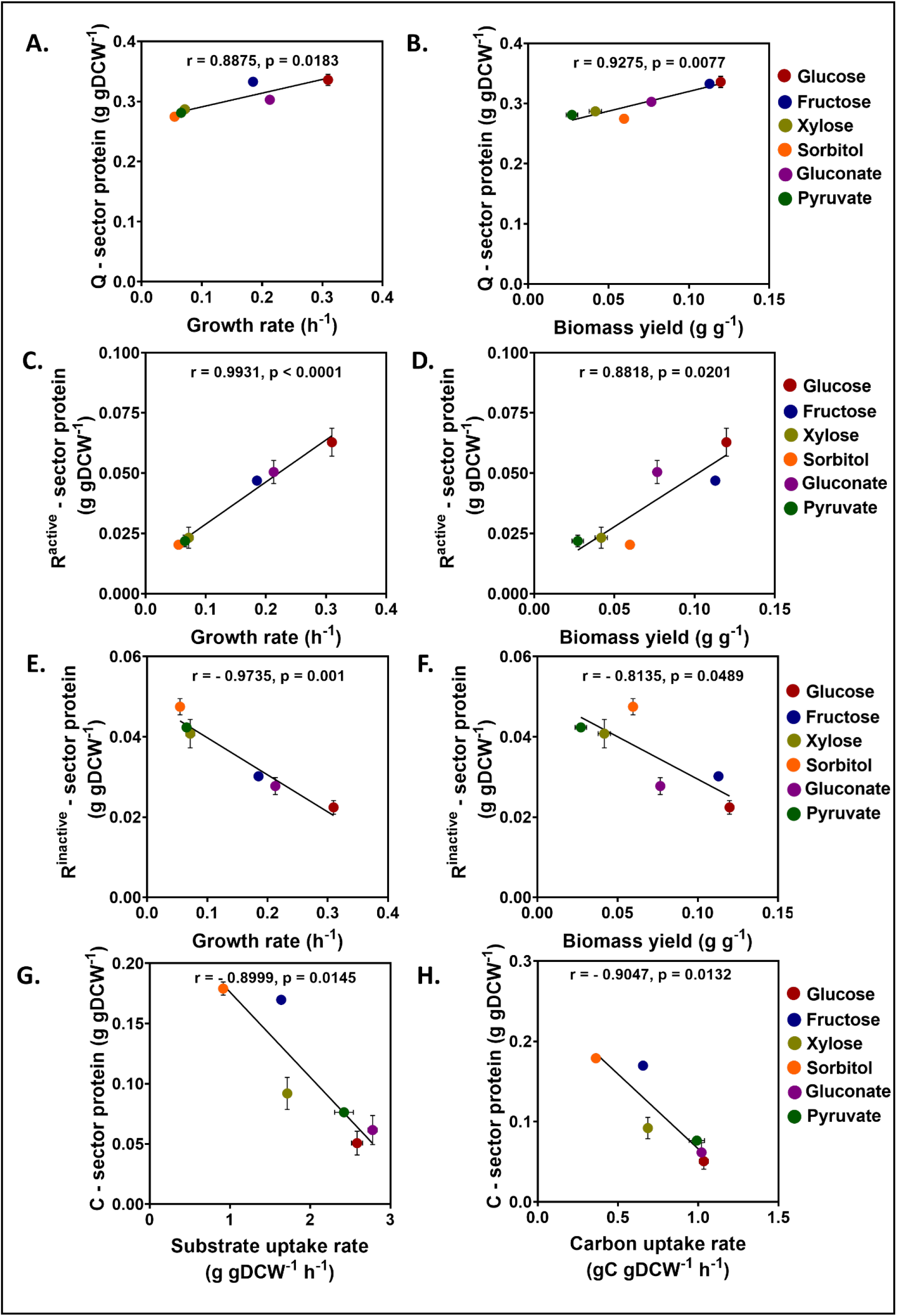
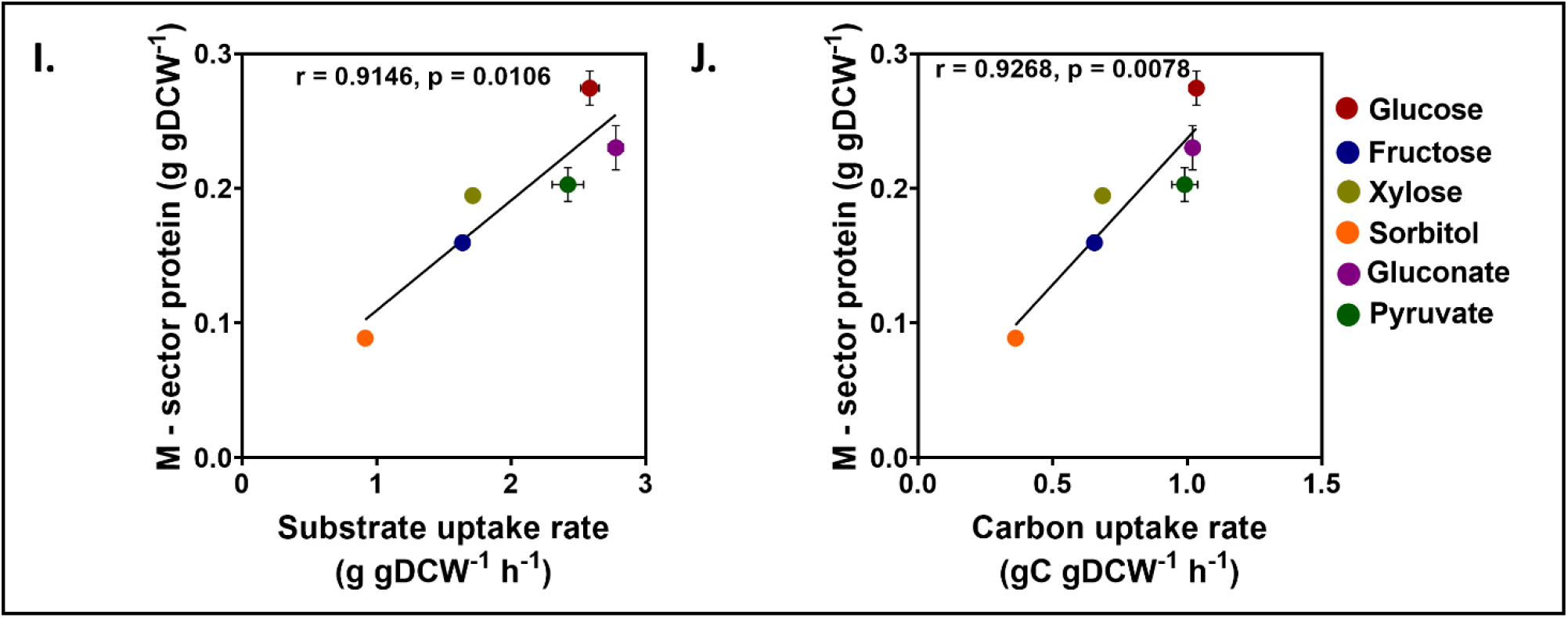
The relationship between absolute protein abundance and physiological parameters. The core proteome abundance with (A) Growth rate and (B) Biomass yield. The active ribosomal proteome abundance with (C) Growth rate and (D) Biomass yield. The inactive ribosomal proteome abundance with (E) Growth rate and (F) Biomass yield. The catabolic proteome abundance with (G) Substrate uptake rate and (H) Carbon uptake rate. The metabolic proteome abundance with (I) Substrate uptake rate and (J) Carbon uptake rate.

Also, the active ribosomal proteome abundance was positively correlated with bacterial proliferation rate and its corresponding biomass yield (Fig. 4 (C) and (D)). On the contrary, inactive ribosomal proteome abundance caused growth retardation, as it was negatively correlated with cellular growth rate and its biomass yield (Fig. 4 (E) and (F)). Additionally, the fraction of active and inactive ribosomal abundances (by weight) were the determinant of cell growth rate and its biomass yield for anaerobic batch fermentation of different substrates.

The substrate influx is a rate limiting step, and the anabolic precursors are derivatives of substrate catabolism. Here, we observed a negative correlation between the substrate influx rate and the corresponding catabolic proteome abundance dedicated for the substrate influx (Fig. 4 (G)). Thus, higher the cellular substrate uptake rate, the lower was the proteome expenditure on its transport and internalization. Similar was the behavior observed on substrate carbon internalization rate. The larger the carbon influx rate in the cell, the lower was the protein expenditure (Fig. 4 (H)). However, the affinity of *E. coli* cells to substrate can also become limiting over here. This is an interesting observation, as the basis of our study is equivalent carbon content. From this result it is imperative that the carbon content is not the determinant of substrate influx in the cell, rather it’s the type of substrate and its uptake mechanism that contribute to cell physiology.

The metabolic sector included biosynthetic proteins contributing to cellular metabolic activities. This sector had lower span in substrates with higher catabolic proteome sector. The higher expense of proteome on substrate internalization, rendered less proteome share for its metabolism, e.g. sorbitol catabolism. As a result, the growth rate was compromised. The biosynthetic sector proteome abundance showed a positive correlation with substrate influx rate and corresponding substrate-carbon influx rate, during anaerobic fermentation of different substrates (Fig. 4 (I) and (J)). This relationship established the substrate requirement for the cellular metabolic functions. An influx of substrate at lower rate would slow down cellular metabolic activities. Consequently, lower metabolic proteins would be channelized for carrying out the necessary cellular biosynthetic activities. As a result,the cell physiology would be compromised under such circumstances.

## Discussions

The cellular ribosomal abundance represented by RNA-to-protein ratio was high during anaerobic fermentation, as compared to aerobic growth conditions (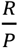 ratio < 0.2) reported in literature for similar growth regime^3,4,25^. The cell cultivation under anoxic (oxygen stress) fermentative (metabolic stress) condition coupled with nutrient limitation would have rendered ribosomal content on higher side. Dai *et al*.^30^ have also reported an increase in cellular ribosome content during hyperosmotic stress in exponentially proliferating *E. coli* cells. The cellular modulation of ribosomal abundance at different growth rates is unsurprisingly rate-limiting in *E. coli* as established previously^4,31^. Considering the anaerobic fermentative catabolism of substrates, the relative proportion of actively translating and non-active ribosomes was critical in determining the *E. coli* growth rate. The fraction of translating ribosomes declined at slower growth rates, which increased the burden on *E. coli* BW25113 cells for optimal behavior. Additionally, under nutrient-limited condition, the bacterial cell doubling time is largely influenced by the time taken by a single ribosome to translate all of its proteins. This in turn, depends on ribosome’s translational capacity and elongation rate determined by the induction of secondary messenger like -ppGpp mediated response in nutrient-limited medium^25,31,32^. In *E. coli*, high ppGpp level is triggered by the enzyme RelA^33^, upon accumulation of de-acylated tRNAs at ribosomal A – site, which enables the cell to adjust its ribosomal content. Also, ppGpp has been reported to inhibit the initiation of DNA replication and its supercoiling near the origin of replication^34,35^, or ppGpp along its co-regulator DksA represses the rRNA synthesis and the gene expression of ribosome affiliated proteins^36,37^. Therefore, the ppGpp overproduction lowers the bacterial growth rate and enhances the stress tolerance in *E. coli* under harsh growth environments^38^. Under such circumstances, the resource re-allocation occur from ribosome synthesis to stress response^38^, mostly metabolic^32^ and catabolic activities under nutrient limited conditions (Fig. 3 (B)). Interestingly, in nutrient limited growth, the proteome efficiency increases along nutrient pathway flow, where substrate uptake and metabolism seem to be less efficient while amino acid biosynthesis and translation utilize proteome efficiently^39^. Our observations regarding higher share of catabolic and metabolic proteome sectors during nutrient- and oxygen-stressed fermentative growth of *E. coli*, are in par with previously reported outcomes. Additionally, the low ATP level during anaerobic fermentation facilitates a decline in rRNA synthesis^40^. Due to the translational feedback inhibition, the amount of free ribosomal protein inhibits their own mRNA, which results in a decline in ribosomal protein synthesis at lower energy levels during anaerobic fermentation. This leads to an increased proportion of inactive ribosomes, along with the active ribosomes translating at a decreased elongation rate. Eventually, *E. coli* cells exhibit slower growth rates and increased ribosomal abundance^30^ during anoxic batch fermentation.

Moving beyond ribosomal dynamics, the nature of substrate metabolism also emerges as a significant determinant of *E. coli*’s physiological response. In anaerobic batch fermentation of glucose, fructose, xylose, sorbitol, gluconate and pyruvate, the nature of substrate metabolism became limiting in providing the requisite energy level for cellular ribosomal activities. Thus, dictating the allocation of cellular resources, impacting ribosomal translation capacity, protein synthesis times, and ultimately, cell doubling rates. In addition, our study identified a modification of the established bacterial growth law, wherein we observed that the relative proportion of active to inactive ribosomes, or the abundance of ribosomal proteins, had a linear impact on bacterial growth rate under conditions of slow proliferation. This can form the basis for studying the cell physiology under different growth conditions and understanding the ribosomal impact on resulting phenotype. A critical bottleneck in cell survivability is substrate influx into cellular environment. This substrate internalization process could be energy consuming, like for xylose influx, or *E. coli* host can possess low affinity for substrate like sorbitol, thereby declining the growth rate. Alternatively, the substrate uptake would happen at comparatively lower pace like fructose, but cellular metabolic wiring would favor substantial proliferation rate. Under such circumstances, the carbon catabolic proteome share would increase, thereby limiting the proteome resource for other cellular activities. Similar observations have been reported for different modes of metabolic limitations^1,14^. The substrates like glucose, gluconate and pyruvate were internalized at higher rate, thus accounting for lower proteome share. However, we couldn’t generalize the relationship between substrate influx rate, catabolic proteome allocation and corresponding growth rate. As these parameters are substrate specific, cells behaved differently under different growth environment. However, if substrate influx happened at higher rate, then lower catabolic proteome share would be allocated for its internalization. Consequently, *E. coli* cell would have higher proteome share for other metabolic activities, thereby improving its performance under different growth conditions.

The core proteome abundance involved in cellular maintenance, increased linearly with *E. coli* growth rate. In order to sustain higher proliferation rate, the bacterial cell expended higher cellular proteome resource on its housekeeping activities. However, the housekeeping proteome fraction remained unaltered across different growth environments. The active ribosomal abundance facilitated bacterial growth rate and its biomass yield. However, opposite was the influence of inactive ribosomal abundance on growth rate and biomass yield. The biosynthetic proteome played a crucial role in maintaining flux across metabolic network operative in *E. coli* cell. These metabolic proteins regulated intermediate bio-conversions necessary for polymerization of macromolecules and cellular energy states. Moreover, the metabolic proteome abundance allocated for such cellular activities, were largely determined by the substrate influx rate. Higher the substrate internalization rate, more would be the biosynthetic proteome requirement to provide precursors for macromolecule polymerization.

## Conclusion

In conclusion, this study elucidates the phenotypic variations resulting from anoxic catabolism of different glycolytic and non-glycolytic substrates by *E. coli*. The critical wiring between host native metabolism and its proteome allocation for different cellular activities, was influenced by the nature of catabolic substrates resulting in distinct phenotypes. The rate-limiting steps like substrate uptake, active ribosomal abundance and translation rate, proteome partitioning, etc. determined the cell specific growth rate under nutrient-limited anoxic condition. These insights advance our understanding about *E. coli* adapting to anaerobic fermentative growth under limited resources. This knowledge of anaerobic metabolism can be explored in understanding the microbial metabolism regulation and efficient management of cellular resources, for sustainable bioprocessing. Future studies can be built upon these findings by exploring ribosomal profiling by improving nutritional capacity, or undertaking the metabolomic and proteomic studies for elaborating the impact of key metabolic reactions and proteome cost associated with bacterial physiology across different medium in an anoxic environment. This can make the findings more pronounce and valuable to scientific communities. Overall, this study recognizes that phenotypic outcomes under different growth conditions are manifestations of altered regulatory controls, including metabolic rewiring and proteome rearrangement. A comprehensive understanding can give a better control over biological conversions and desired phenotypes, thereby minimizing the cost associated with product driven bioconversions.

## Conflict of Interest

The authors declare no competing interests.

## Notes

### Competing Interest Statement

The authors have declared no competing interest.

