## Supplementary file for "The catabolic nature of fermentative substrates influence proteomic rewiring in *Escherichia coli* under anoxic growth"

### S.1 Relationship between active ribosomal fraction and bacterial growth rate

In nutrient-limited anaerobic batch fermentation of glucose, fructose, xylose, sorbitol, gluconate, and pyruvate; the *E. coli* BW25113 cells exhibited different proliferation rates. As established previously, a decline in actively translating ribosomes was linked to a compromised growth rate in *E.coli*[1]. Thus, we consolidated the data from the literature[1,2], depicting the fraction of active ribosomes in *E. coli* at different growth rates. We did curve fitting and observed the Michaelis-Menten type of relationship between active ribosomal fraction and growth rate (Fig. S1. A). Both the parameters were related by the equation:

$$f_{active} = \frac{f_{active}^{max} \times \mu}{K_g + \mu} \qquad \text{(S.1)}$$

where $f_{active}^{max} = 1.073$ is the maximum fraction of active ribosomes in *E.coli* cells and $K_g = 0.1412$ h$^{-1}$ is bacterial growth rate corresponding to an active ribosomal fraction when it is exactly half of its maximum value and $\mu$ (h$^{-1}$) corresponds to respective bacterial growth rates.

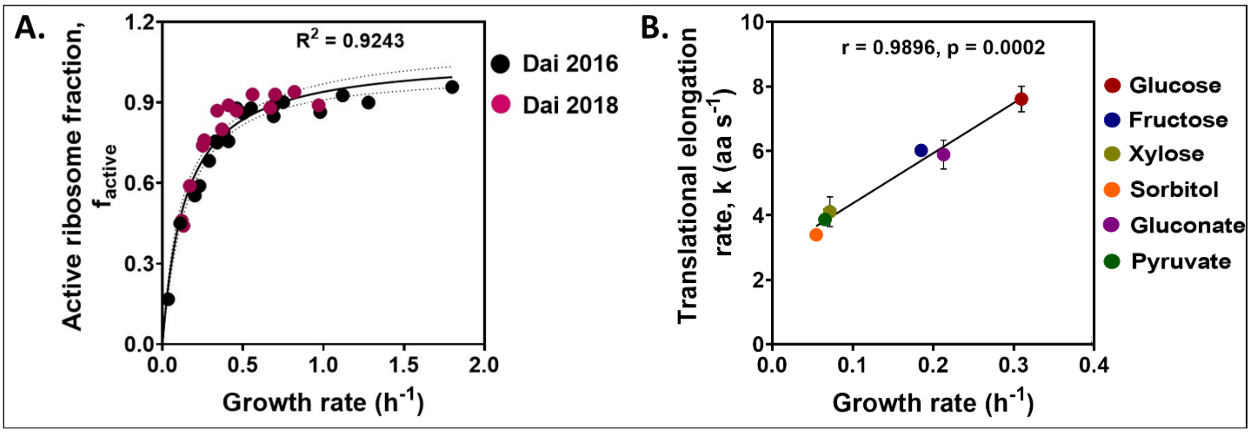


Fig. S1: (A) The relationship between fraction of *E. coli* active ribosomes and growth rate[1,2]. (B) The correlation between translation elongation rate and *E. coli* growth rate during anaerobic fermentative metabolism of different substrates.

The translation elongation rate of active ribosomes is an important parameter influencing cell proliferation rates. The relationship between active ribosomal fraction and translation elongation rate has been defined as[1]:

$$f_{active} = \frac{\mu \times \sigma'}{k \times \frac{R}{P}} \quad \text{(S.2)}$$

where *k* is the translation elongation rate (amino acid per second), $\sigma' \equiv \frac{m_{rRNA}}{0.86 \times m_{aa}} \approx 1.56 \times 10^4$ is a dimensionless constant, the $\frac{R}{P}$ represents ribosomal abundance. We equated equations S1 and S2, and deduced a formula for determining the translation elongation rate given as

$$k = \frac{(K_g + \mu) \times \sigma'}{f_{active}^{max} \times \frac{R}{P}} \quad \text{(S.3)}$$

Thus, the translation elongation rate is directly correlated with bacterial proliferation rate. The linearity between theoretical translation elongation rate and *E. coli* BW25113 growth rate during anaerobic batch fermentation of different glycolytic and non-glycolytic substrates is depicted in figure S1 (B). To further verify our theoretical considerations, we calculated the translation elongation rate using the carbon-limited ribosomal pool model defined by Li *et al.* (2018). The values predicted by using equation S3 were same as those predicted by ribosomal pool model[3], thus we considered our values to be within acceptable limits. However ribosomal profiling experiments can be carried out in future to validate the theoretical predictions.

### S.2 The influence of ribosomal proteome share on *E. coli* BW25113 growth rate

The relationship between cellular ribosome content and ribosome affiliated proteome fraction was defined as[1]:

$$\emptyset_R = \frac{r}{\sigma} \quad \text{(S.4)}$$

where $\emptyset_{Rb}$ is the proteome fraction of ribosomes, *r* is the cellular ribosome content (RNA-to-protein ratio, $\frac{R}{P}$) and $\sigma = \frac{m_{rRNA}}{0.86 \times m_{Rb}} \approx 2.1$ (since $m_{rRNA}$ = 1,479,384 is mass of ribosomal RNA and $m_{Rb}$ = 806,960 is mass of ribosomal proteins contained in a single ribosome)[1]. The total ribosomal proteome can be split in to its active and inactive fraction as:

$$\emptyset_R^{active} = f_{active} \times \emptyset_R \quad \text{(S.5)}$$

$$\emptyset_R^{inactive} = \emptyset_R - \emptyset_R^{active} \quad \text{(S.6)}$$

the active and inactive portion of ribosomal proteome are represented by $\emptyset_R^{active}$ and $\emptyset_R^{inactive}$ respectively.

Scott et al. (2010) have defined a linear relationship between cellular ribosome content ($r$) and growth rate ($\mu$) as:

$$r = r_0 + \frac{\mu}{k_t} \quad \text{(S.7)}$$

where $r_0$ is the vertical intercept and $k_t$ is the inverse of the slope known as the "translation capacity" of organism. However, Dai et al. (2016) have modified the equation S7 by introducing a constant $\sigma$, which can be re-written in terms of ribosomal proteome abundance as:

$$\emptyset_R = \emptyset_{R,0} + \frac{\mu}{k_t'} \quad \text{(S.8)}$$

Where $\emptyset_{Rb,0} = \frac{r_0}{\sigma}$ is the basal ribosomal proteome and $k_t' = k_t \times \sigma$. The equations S.7 and S.8 were applied under a constant translation elongation rate. However, during the existence of active and inactive ribosomal fractions within a cell, the equation S8 was modified as:

$$\emptyset_R = \emptyset_R^{inactive} + \frac{\mu}{\gamma} \quad \text{(S.9)}$$

The basal ribosomal proteome fraction can be attributed as inactive ribosomal proteome, owing to its non-significant contribution in determining the cell proliferation rate and $\frac{1}{\gamma}$ is the time taken by one active ribosome to synthesize all ribosomal proteins[1].

### S.3 Defining the model parameters for cellular proteome partitioning

The constrained allocation flux balance analysis[4] was used to allocate proteome share into four coarse-grained sectors, namely, ribosomal sector (R-sector), catabolic sector (C-sector), metabolic sector (M-sector), and housekeeping sector (Q-sector) respectively. The proteome mass fractions for each sector were represented by $\emptyset_X$, where X ∈ (R, C, M, Q). These proteome fractions adjust with bacterial growth rate under different growth environments and mathematically add up to unity, as:

$$\emptyset_R + \emptyset_C + \emptyset_M + \emptyset_Q = 1 \quad \text{(S.10)}$$

Rearranging,

$$\emptyset_R + \emptyset_C + \emptyset_M = 1 - \emptyset_Q \quad \text{(S.11)}$$

The term on right-hand-side is fixed and the variable proteome allocation would happen among other three functional sectors. Based on bacterial growth law[5], each of the condition-dependent terms in equation S.11 is of the form[4]:

$$\emptyset_X = \emptyset_{X,0} + \Delta\emptyset_X \tag{S.12}$$

where $\emptyset_{X,0}$ is a constant offset value and its minimum value can be zero, and X ∈ (R, C, M). The $\Delta\emptyset_X$ is characterized by the driving fluxes in each sector. Now extending the equation S.12 relationship to each variable proteome sector, we have:

1. The R-sector ($\emptyset_R$): Under nutrient-limited bacterial growth, $\emptyset_R$ is linearly related with growth rate μ as[4–6]:

$$\emptyset_R = \emptyset_{R,0} + w_R \times \mu \tag{S.13}$$

   where $\emptyset_{R,0}$ represents extrapolated basal ribosomal protein at no growth and is strain dependent, and phenomenologically $w_R$ is the proteome fraction allocated to ribosomal protein per unit growth rate[4].

   In this study, we have equation S.9 similar to equation S.12. On comparing these two equations, the extrapolated basal ribosomal proteome ($\emptyset_{R,0}$), can be considered equivalent to the inactive ribosomal proteome fraction ($\emptyset_R^{inactive}$). Moreover, the phenomenological parameter $w_R$ analogs to ribosomal protein synthesis time ($\frac{1}{\gamma}$). We have already determined the values of $\emptyset_R^{inactive}$ and $\frac{1}{\gamma}$ for *E. coli* BW25113 cells growing anaerobically in glucose, fructose, xylose, sorbitol, gluconate and pyruvate. These parameters for individual growth conditions, were used as input while simulating the model.
2. The M-sector: Mori et al. (2016) have defined a linear relationship between cell proliferation rate and enzyme abundance[7,8] under carbon limited growth. Also, considering linear dependence between growth rate and biosynthetic fluxes as:

$$\emptyset_M = \emptyset_{M,0} + \textstyle\sum_{i\,\epsilon\,M} w_i\,|v_i| \tag{S.14}$$

   where $\emptyset_{M,0}$ is a fixed offset (minimum can be zero) and $w_i$ is the proteome fraction allocated in enzyme $M_i$ per unit of reaction *i*, that was assumed to take same value for all the reactions as $w_M = 8.3 \times 10^{-4}$ g h mmol[-1][4].
3. The C-sector: Considering the experimental findings of bacterial growth on a single carbon source[7,8], the authors Mori et al. (2016) have considered linear relationship between carbon catabolism proteome ($\emptyset_C$) and carbon intake flux ($v_C$) as:

$$\emptyset_C = \emptyset_{C,0} + w_C \times |v_C| \tag{S.15}$$

where $\emptyset_{C,0}$ corresponds to offset (growth independent), and $w_C$ is C-sector proteome per unit carbon influx[4]. For our study, we considered the catabolic and metabolic offsets to be at their minimal value.

The above parameters were used to simulate genome-scale metabolic model *i*JO1366 (with necessary modifications for resembling the genetic makeup of *E. coli* BW25113), for partitioning the cellular proteome into four coarse-grained functional sectors.
